# Integrative Transcriptomic and Experimental Analysis Reveals Stage-Associated Molecular Changes and Context-Dependent Expression Responses of CDKN1A and c-MYC in NAFLD Progression

**DOI:** 10.64898/2026.11.26.769377

**Authors:** Zahra Moayedfard, Negar Azarpira, Shima Parsa, Kimia Falamarzi, Farnaz Sani, Ayeh Bolouki

## Abstract

**Background and Aims:** Non-alcoholic fatty liver disease (NAFLD) progression involves complex molecular changes across disease stages, but the regulatory programs underlying these transitions remain incompletely understood. This study aimed to characterize stage-associated molecular changes and investigate the expression responses associated with CDKN1A and c-MYC overexpression.

**Materials and Methods:** Transcriptomic data from 114 human liver samples across five NAFLD stages were analyzed using differential gene expression, weighted gene co-expression network analysis (WGCNA), and protein–protein interaction (PPI) network analysis. CDKN1A and c-MYC were further investigated through overexpression experiments in normal and fatty-acid-treated HepG2 cells, followed by quantitative PCR analysis of selected genes.

**Results:** A total of 3,663 differentially expressed genes (DEGs) were identified across four consecutive disease-stage transitions, with the largest transcriptional changes occurring between NASH and mild fibrosis. WGCNA identified stage-associated co-expression modules enriched in inflammatory, cell-cycle, and tissue-remodeling pathways. PPI analysis revealed stage-dependent changes in interaction networks involving immune signaling, transcriptional regulation, and metabolic processes. In HepG2 cells, CDKN1A overexpression was associated with changes in inflammatory and TNF-related gene expression, whereas c-MYC overexpression was associated with increased expression of cell-cycle-related genes. These responses varied between normal and steatotic cells and across individual genes.

**Conclusions:** NAFLD progression is characterized by stage-associated transcriptional and network remodeling. CDKN1A and c-MYC overexpression are associated with distinct gene-expression responses related to inflammation and cell-cycle regulation, highlighting their potential relevance to hepatocyte responses under metabolic stress.

## INTRODUCTION

Non-alcoholic fatty liver disease (NAFLD) is one of the most common chronic liver disorders worldwide, encompassing a spectrum of conditions ranging from simple steatosis to non-alcoholic steatohepatitis (NASH), progressive fibrosis, and, in some patients, hepatocellular carcinoma. Advanced liver fibrosis is a major predictor of liver-related complications and mortality, highlighting the importance of understanding the molecular mechanisms underlying disease progression. Despite increasing recognition of NAFLD as a major public health concern, therapeutic options remain limited, particularly for patients with progressive liver disease(Powell, Wong, & Rinella, 2021; Solleiro-Villavicencio, Viurcos-Sanabria, Aguayo-Guerrero, Pineda-Pérez, & Méndez-García, 2025; Younossi et al., 2023).

Cellular senescence and stress-response pathways are increasingly recognized as important contributors to NAFLD progression. Under metabolic and lipotoxic stress, hepatocytes may develop senescence-associated secretory phenotypes (SASP), characterized by the release of inflammatory mediators that can influence the hepatic microenvironment and contribute to tissue remodeling. Cyclin-dependent kinase inhibitor 1A (CDKN1A, also known as p21), a major regulator of cell-cycle arrest and a downstream effector of p53, plays an important role in cellular stress responses. Although transient CDKN1A activation may contribute to protective cell-cycle arrest, sustained expression has been associated with cellular senescence, inflammation, and fibrogenic processes (Shan et al., 2024; D. S. Umbaugh, Nguyen, Smith, Ramachandran, & Jaeschke, 2024; Xing & Zhu, 2025).

Another important regulator of cellular growth and metabolism is cellular MYC proto-oncogene (c-MYC), a transcription factor involved in cell-cycle progression, metabolic regulation, and cellular proliferation. Altered c-MYC expression has been implicated in hepatic metabolic dysfunction, steatohepatitis, and liver tumor development(Guo et al., 2021; Jia, Che, Cigliano, Wang, Peitta, Tao, Zhong, Ribback, Evert, & Chen, 2020). Given the distinct biological functions of CDKN1A and c-MYC, changes in their expression may contribute to different aspects of hepatocyte responses during NAFLD progression. However, their stage-associated expression patterns and potential roles in inflammatory and cell-cycle-related responses under metabolic stress remain incompletely understood.

In this study, we integrated transcriptomic and network-based analyses of human liver samples representing different stages of NAFLD progression to identify stage-associated molecular changes and potential regulatory genes. We further investigated the effects of CDKN1A and c-MYC overexpression, individually and in combination, in normal and fatty-acid-treated HepG2 cells. The expression of selected genes involved in inflammation, immune regulation, metabolism, cell-cycle control, and tumor necrosis factor (TNF)-related signaling was examined to explore how these two regulators are associated with hepatocyte transcriptional responses under different metabolic conditions.

## METHODS

### Dataset Selection and Sample Classification

Human liver gene expression datasets (GSE48452, GSE49541, and GSE63067), comprising 114 samples across five disease stages, were retrieved from the Gene Expression Omnibus (GEO) database(Ahrens et al., 2013; Frades et al., 2015; Murphy et al., 2013). The samples were classified as normal (n = 8), steatosis (n = 8), NASH (n = 26), mild fibrosis (F1–F2; n = 40), and advanced fibrosis (F3–F4; n = 32). The original studies had received ethical approval, as documented in their respective publications.

### Data Preprocessing and Batch Correction

Raw expression data from the three GEO datasets (GSE48452, GSE49541, and GSE63067), generated using the same microarray platform, were combined into a single expression dataset. Quality assessment, normexp background correction, quantile normalization, and log₂ transformation were performed using the GEOquery, affy, oligo, and limma packages in R(Carvalho & Irizarry, 2010; Davis & Meltzer, 2007; Gautier, Cope, Bolstad, & Irizarry, 2004; M. E. Ritchie et al., 2015). To account for inter-dataset technical variation while preserving biological differences associated with disease progression, ComBat batch correction was performed once on the combined dataset using the sva package in R(Leek, Johnson, Parker, Jaffe, & Storey, 2012). GEO dataset identity was specified as the batch variable, and disease stage was included as a biological covariate(Matthew E. Ritchie et al., 2006).

### Partial Least Squares Discriminant Analysis

Partial least squares discriminant analysis (PLS-DA) was performed in R using the ropls package to investigate the separation of liver samples according to disease stage(Thévenot, Roux, Xu, Ezan, & Junot, 2015). The normalized gene expression matrix was used as the predictor dataset, and disease-stage classifications were used as the categorical response variable. Two separate PLS-DA models were constructed: the first compared normal, steatosis, and NASH samples, while the second compared NASH, mild fibrosis (F1–F2), and advanced fibrosis (F3–F4) samples. The default component-selection settings of the ropls package were used. Model validity was assessed using permutation testing with 100 permutations. Score plots were generated to visualize sample distributions according to disease stage.

### Differential Expression Analysis

Differential gene expression analysis was performed using the limma package in R. Four separate pairwise comparisons were conducted using the batch-corrected expression dataset to investigate transcriptional changes across consecutive stages of NAFLD progression: steatosis versus normal, NASH versus steatosis, mild fibrosis (F1–F2) versus NASH, and advanced fibrosis (F3–F4) versus mild fibrosis (F1–F2). Genes with an absolute log₂ fold change greater than 0.5 and a Benjamini–Hochberg-adjusted P-value less than 0.05 were considered differentially expressed(M. E. Ritchie et al., 2015).

### Weighted Gene Co-expression Network Analysis

Weighted gene co-expression network analysis (WGCNA) was performed using the WGCNA R package to identify gene co-expression modules associated with NAFLD progression(Langfelder & Horvath, 2008). The 5,000 most variably expressed genes across all samples were selected for network construction. Samples were hierarchically clustered to identify potential outliers.

Separate unsigned co-expression networks were constructed for each of the four consecutive disease-stage comparisons: normal versus steatosis, steatosis versus NASH, NASH versus mild fibrosis (F1–F2), and mild fibrosis (F1–F2) versus advanced fibrosis (F3–F4). Only samples belonging to the two stages under comparison were included in each network analysis. The analyses were performed using the batch-corrected expression matrix obtained after ComBat correction.

Pairwise Pearson correlation coefficients were calculated between genes, and an unsigned adjacency matrix was generated using soft-thresholding powers (β) of 5, 5, 9, and 15 for the four respective comparisons. The adjacency matrix was subsequently transformed into a topological overlap matrix (TOM) to assess network interconnectedness. Gene modules were identified through hierarchical clustering and the dynamic tree cut algorithm, with a minimum module size of 30 genes. Module eigengenes (MEs), representing the first principal component of each module, were calculated and correlated with clinical traits.

Clinical traits included disease-stage status and numerical fibrosis scores obtained from GEO sample annotation files for samples in the normal, steatosis, and NASH groups. For samples in the fibrosis groups, disease severity was represented by the predefined categories of mild fibrosis (F1–F2) and advanced fibrosis (F3–F4).

Gene significance (GS) was defined as the correlation between individual gene expression and clinical traits, whereas module membership (MM) represented the correlation between individual gene expression and its corresponding module eigengene. Module–trait associations with Benjamini–Hochberg-adjusted P-values below 0.05 were considered statistically significant. Genes with |GS| > 0.2 and |MM| > 0.8 within significant modules were designated as hub genes.

### Protein-Protein Interaction Network Analysis

To explore functional interactions among proteins encoded by DEGs, protein-protein interaction (PPI) networks were constructed. Gene symbols of interest were mapped to the STRING database (v11.5) (Szklarczyk et al., 2019) to retrieve experimentally validated and high-confidence predicted interactions, with a combined score threshold > 0.4 to retain medium-and high-confidence interactions.

PPI networks were visualized and analyzed using Cytoscape (v3.9.1) (Shannon et al., 2003). To identify densely connected clusters and potential functional modules, the MCODE plugin (Bader & Hogue, 2003) was applied using default parameters (degree cutoff = 2, node score cutoff = 0.2, k-core = 2, max depth = 100). Clusters with the highest MCODE scores were considered key functional modules and subjected to functional enrichment analysis using clusterProfiler, to elucidate pathways and processes potentially driving NAFLD progression and fibrosis.

Additionally, hub genes within the PPI networks were identified based on topological features such as degree, betweenness, and closeness centrality, calculated using the cytoHubba plugin (Chin et al., 2014).Hub genes were defined as those ranking in the top 5% for at least two centrality measures, reflecting their potential regulatory significance within the network (Bader & Hogue, 2003; Chin et al., 2014; Shannon et al., 2003; Szklarczyk et al., 2019).

### Functional Enrichment Analysis

To investigate the biological relevance of differentially expressed genes (DEGs), WGCNA modules, and PPI network clusters, functional enrichment analysis was performed using the clusterProfiler R package(G. Yu, Wang, Han, & He, 2012). Gene Ontology (GO) enrichment analysis was conducted across three categories: Biological Process (BP), Molecular Function (MF), and Cellular Component (CC). In addition, pathway enrichment analyses were performed using the Kyoto Encyclopedia of Genes and Genomes (KEGG), Reactome, and WikiPathways databases to identify relevant signaling and metabolic pathways.

All enrichment results were corrected for multiple testing using the Benjamini–Hochberg method, and terms with an adjusted P-value (false discovery rate, FDR) below 0.05 were considered statistically significant. Functional enrichment analyses were performed separately for DEG lists, genes belonging to significant WGCNA modules, and genes within selected PPI network clusters to identify biological processes and pathways associated with NAFLD progression and fibrosis development.

### Cell Culture, Steatosis Induction, and Experimental Design

HepG2 cells were cultured in Dulbecco’s modified Eagle medium (DMEM) supplemented with 10% fetal bovine serum (FBS) at 37°C in a humidified atmosphere containing 5% CO₂. Steatosis was induced by treating cells with oleic acid (1 mM) and palmitic acid (0.5 mM) at a 2:1 ratio for 24 hours. Intracellular triglyceride levels were quantified to assess lipid accumulation, and steatosis was further validated by evaluating the expression of lipogenic markers, including SREBP-1c, FASN, CD36, and PLIN2(Ricchi et al., 2009). Following steatosis induction, cells were transfected and maintained for an additional 72 hours before harvest.

### Plasmid Construction and Transfection

The full-length coding sequences of human *CDKN1A* and *MYC*, each fused to a Myc epitope tag, were subcloned into the bicistronic pIRES2-EGFP vector (Clontech/Takara). In this vector, the gene of interest and EGFP are expressed as separate proteins through an internal ribosome entry site; therefore, EGFP protein expression served as an independent marker of transfection efficiency. The constructs were verified by Sanger sequencing.

HepG2 cells at approximately 80% confluence were transfected with 2 µg of total plasmid DNA using Lipofectamine with PLUS Reagent (Thermo Fisher Scientific). Following a 4-hour transfection period, cells were maintained for an additional 72 hours before harvest. Five experimental conditions were evaluated under both normal and steatotic conditions: untransfected control, empty vector, *CDKN1A* overexpression, *MYC* overexpression, and *CDKN1A/MYC* co-transfection. For co-transfection, 1 µg of the *CDKN1A* plasmid and 1 µg of the *MYC* plasmid were used (1:1 ratio). Total plasmid DNA was balanced across groups using empty vector. Each experimental group included five biological replicates. Transfection efficiency was monitored by EGFP protein expression, and expression of Myc-tagged CDKN1A and c-Myc was confirmed by Western blotting using an anti-Myc-tag antibody.

### Western Blot and qPCR Analysis

Protein expression was assessed by Western blotting to confirm successful transfection. EGFP was detected as an independent marker of transfection, while Myc-tagged CDKN1A and c-MYC proteins were detected using an anti-Myc epitope-tag antibody. Proteins were extracted using RIPA buffer, and membranes were incubated overnight at 4°C with the anti-Myc antibody (1:500 dilution). Protein bands were visualized using enhanced chemiluminescence.

Total RNA was isolated using TRIzol reagent and reverse-transcribed into complementary DNA. Quantitative PCR was performed to assess the expression of 24 selected genes involved in inflammatory signaling, immune responses, TNF-related pathways, cell-cycle regulation, metabolism, and transcriptional regulation. Each of the five biological replicates per experimental group was analyzed in triplicate technical reactions(Bustin et al., 2009).

### Statistical Analysis

Statistical analyses were performed using GraphPad Prism (GraphPad Software, USA). Quantitative PCR experiments were conducted using five independent biological replicates per experimental group, with each biological sample analyzed in triplicate technical reactions. Technical replicates were averaged before statistical analysis. Relative gene expression was calculated using the 2⁻ΔΔCt method, with B2M as the reference gene. For each gene, CDKN1A overexpression, c-MYC overexpression, and CDKN1A/c-MYC co-transfection groups were compared with the corresponding empty-vector control. These comparisons were performed separately in normal and steatotic HepG2 cells. Data are presented as mean ± SEM. Statistical significance was defined as P < 0.05.

## RESULTS

### Stage-Specific Transcriptomic Dynamics Reveal Progressive Molecular Remodeling

Analysis of three microarray datasets comprising 114 samples across five NAFLD stages identified 3,663 dDEGs using the thresholds |log₂FC| > 0.5 and Benjamini–Hochberg-adjusted P < 0.05. PLS-DA demonstrated separation of liver samples according to disease stage (Figure 1A–B), indicating distinct transcriptomic profiles across NAFLD progression.

**Figure 1.**
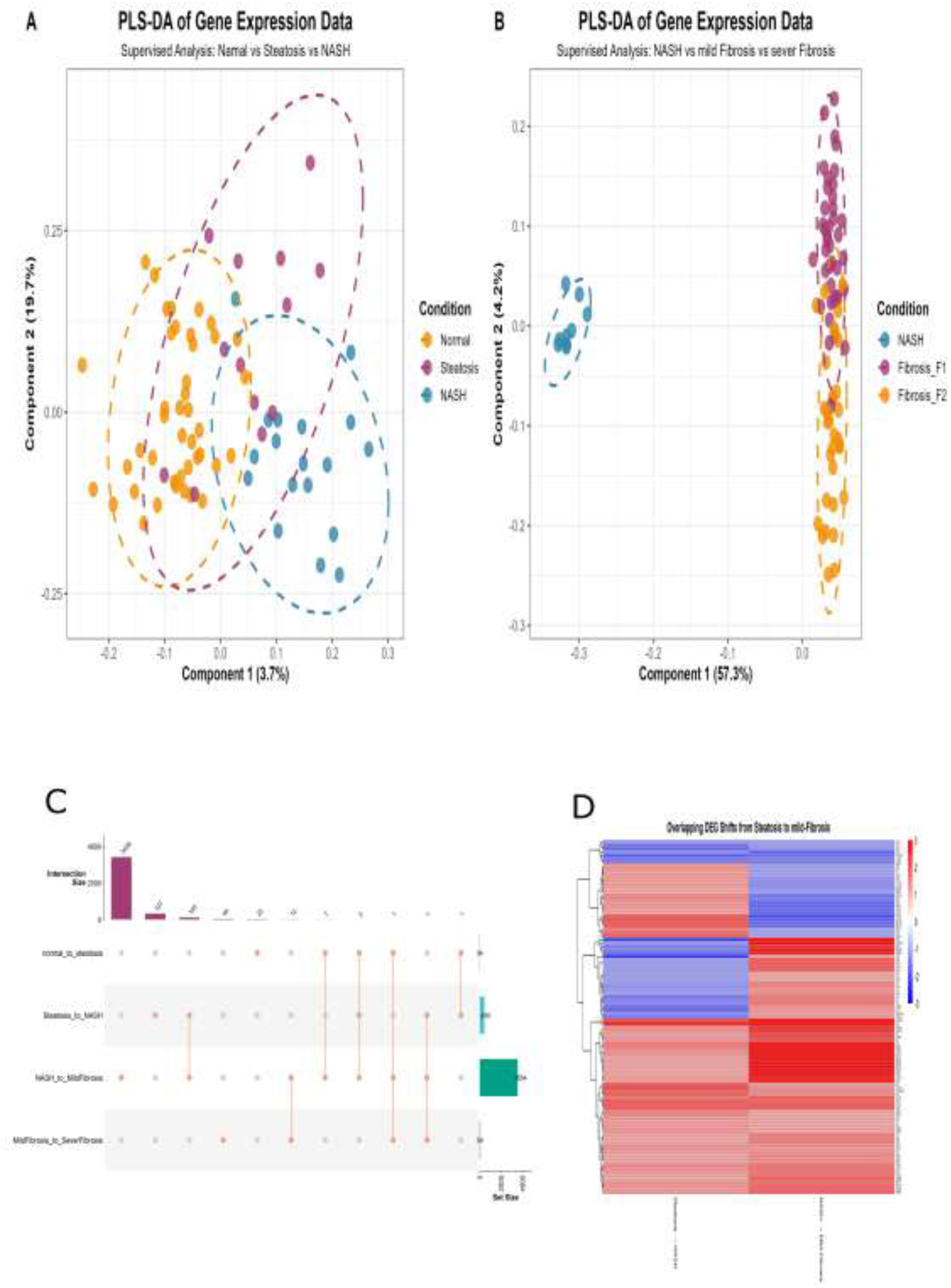
Stage-specific transcriptomic remodeling across NAFLD progression. (A) Partial least squares discriminant analysis (PLS-DA) score plot showing the distribution of liver samples among the Normal, Steatosis, and NASH groups. (B) PLS-DA score plot showing the distribution of liver samples among the NASH, Mild fibrosis (F1–F2), and Advanced fibrosis (F3–F4) groups. Each point represents an individual sample. Percentages on the axes indicate the proportion of variance explained by the corresponding component. Dashed ellipses represent 95% confidence regions for each group. (C) UpSet plot showing differentially expressed genes (DEGs; |log2FC| > 0.5, adjusted P < 0.05) across the four disease-stage transitions. Vertical bars indicate intersection sizes, while horizontal bars indicate the total number of DEGs in each transition, showing predominantly transition-specific expression changes with limited overlap. (D) Heatmap of the 105 DEGs shared between the Steatosis-to-NASH and NASH-to-Mild fibrosis transitions, hierarchically clustered according to log2 fold change. Red and blue indicate upregulation and downregulation, respectively. Among these genes, 46 show inverse regulation between the two transitions, suggesting dynamic transcriptional remodeling.

The number of DEGs varied substantially across consecutive disease-stage transitions. Normal-to-steatosis progression involved 34 DEGs, indicating relatively limited early transcriptional changes. The steatosis-to-NASH transition involved 368 DEGs, consistent with more extensive transcriptional remodeling. The NASH-to-mild fibrosis (F1–F2) transition showed the largest number of DEGs (3,208), involving changes in extracellular matrix organization, immune-related processes, and hepatocyte functions. In contrast, the mild-to-advanced fibrosis (F3–F4) transition involved 52 DEGs, suggesting a more restricted transcriptional response during advanced fibrosis progression (Figure 1C, Table 1, Supplementary File 1).

**Table 1.** Top 10 DEGs ranked by logFC.

|  | **Steatosis vs Normal** |  | **NASH vs Steatosis** |  | **Mild fibrosis vs NASH** |  | **Advanced fibrosis vs Mild fibrosis** |  |
|---|---|---|---|---|---|---|---|---|
| **Rank** | **Up** | **Down** | **Up** | **Down** | **Up** | **Down** | **Up** | **Down** |
| **1** | B3GAT1 | CXCL10 | MIS18BP1 | GSTT1 | EGR1 | CYP7A1 | CHI3L1 | DHRS2 |
| **2** | NT5E | CXCL9 | SLC22A10 | PLEK | VMP1 | ZMAT3 | EPCAM | MT1M |
| **3** | IL1RL2 | CCL20 | HBA2-1 | G6PC | MALAT1 | MALAT1 | CD24 | CYP2C19 |
| **4** | RORA | CDKN1A | TOMM6 | COMMD9 | NEAT1 | WASL | EFEMP1 |  |
| **5** |  |  | CCL18 | MYC | GPAA1 | MALAT1 | RAB12 | CXCL6 |
| **6** |  |  | LGALS3 | PRKAR2B | ZNF160 | APOB | HECTD1 | CHI3L1 |
| **7** |  |  | CD19 | GDF15 | TGDS | SOCS3 | ATP6V0E1 | COL1A2 |
| **8** |  |  | CCR1 | SNORA27 | PPP1R3B | VMP1 | CDC42BPA | AKR1B10 |
| **9** |  |  | CXCR4 | LHFP | DDTL | CCNL1 | CEBPA | ITGBL1 |
| **10** |  |  | CD53 | AIF1 | DACH2 | NAMPT | SMC3 | EPHA3 |

Several genes exhibited stage-dependent expression patterns. CDKN1A was initially downregulated in steatosis and subsequently upregulated through mild fibrosis, suggesting potential involvement in cell-cycle regulation and fibrotic remodeling. CCL3 showed decreased expression in steatosis followed by increased expression in NASH and mild fibrosis, indicating stage-dependent changes in inflammatory signaling. Among the 105 DEGs shared between the steatosis-to-NASH and NASH-to-mild fibrosis transitions, 46 displayed inverse expression patterns, being upregulated in one transition and downregulated in the other. These genes were predominantly associated with inflammatory and nuclear factor kappa B (NF-κB)-related pathways, suggesting dynamic transcriptional remodeling during disease progression (Figure 1D).

### Gene Set Enrichment Analysis Reveals Sequential Pathway Dysregulation

Gene set enrichment analysis identified stage-specific pathway alterations. In steatosis versus normal, immune/chemokine pathways were suppressed—including G protein-coupled receptor signaling, chemokine-mediated signaling, and mononuclear cell migration (enrichment score = −0.81, adjusted p < 0.05, Figure 2A)—driven by decreased CXCR4, CCR1, CCL18, CCL20, CXCL9, and CXCL10. Conversely, cytokine production and carbohydrate metabolism were upregulated, reflecting metabolic adaptation with impaired immune surveillance (Supplementary File 2). Steatosis→NASH transition showed upregulated hepatocyte proliferation, lipid metabolism, and negative angiogenesis regulation (enrichment score > 0.6, adjusted p < 0.05), while immune activation pathways—inflammatory response, interferon-gamma production, interleukin-10 signaling, endocytosis—were downregulated (Figure 2B, Supplementary File 2).

**Fig. 2.**
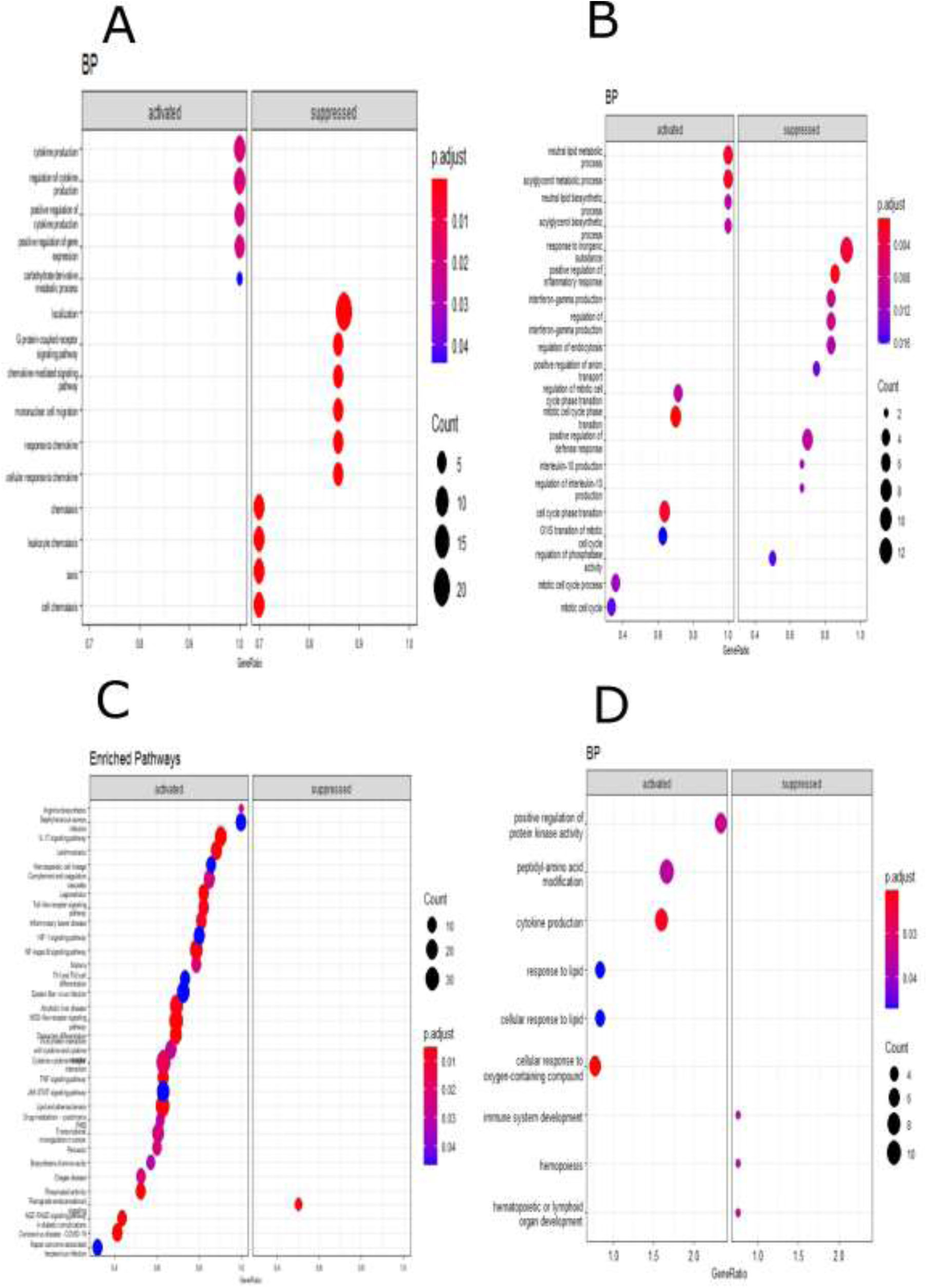
Stage-specific pathway dysregulation across NAFLD transitions identified by gene set enrichment analysis. Enriched terms are shown separately for activated and suppressed gene sets in each disease transition: **a** normal→steatosis (Gene Ontology biological process, BP), **b** steatosis→NASH (BP), **c** NASH→mild fibrosis (KEGG pathways), and **d** mild→advanced fibrosis (BP). Dot size represents the number of genes associated with each term and dot colour represents the adjusted *P* value; the *x* axis indicates the gene ratio. Early steatosis is characterised by suppression of chemokine signalling and immune-cell migration pathways alongside activation of cytokine production and carbohydrate metabolism, whereas later transitions show progressive activation of inflammatory, metabolic, and pro-fibrotic programmes

NASH→mild fibrosis involved extensive remodeling with upregulated stress responses, metabolism, cell differentiation, immune activation, and pro-fibrotic pathways (lipid/atherosclerosis, NOD-like receptor, TNF, NF-κB, advanced glycation end product-receptor for advanced glycation end product signaling). Concurrent proteolytic and endocannabinoid pathway downregulation indicated shift toward extracellular matrix deposition and tissue remodeling (Figure 2C, Supplementary File 2). Advanced fibrosis displayed activated responses to oxygen-containing compounds, cytokine production, protein kinase activity, and peptidyl-amino acid modification, while immune development, hemopoiesis, and lymphoid organ development pathways were suppressed (|enrichment score| > 0.6, adjusted p < 0.05, Figure 2D, Supplementary File 2).

### WGCNA Identifies Stage-Associated Co-Expression Modules

Transition-specific WGCNA analyses were performed using samples from the two disease stages included in each comparison. Four modules showed significant associations with disease-stage or clinical traits across the first three transitions (Supplementary File 3).

In the normal-to-steatosis comparison, the Red module was negatively correlated with steatosis status and numerical fibrosis score (|r| > 0.25, Benjamini–Hochberg-adjusted P < 0.05; Figure 3A). This module was enriched in 5S ribosomal RNA transcription and actin filament depolymerization pathways (Figure 3B). The Yellow module was positively associated with steatosis (|r| > 0.25, adjusted P < 0.05; Figure 3A) and showed enrichment in cytokine production regulation and complement and coagulation cascades (Figure 3C).

**Fig. 3.**
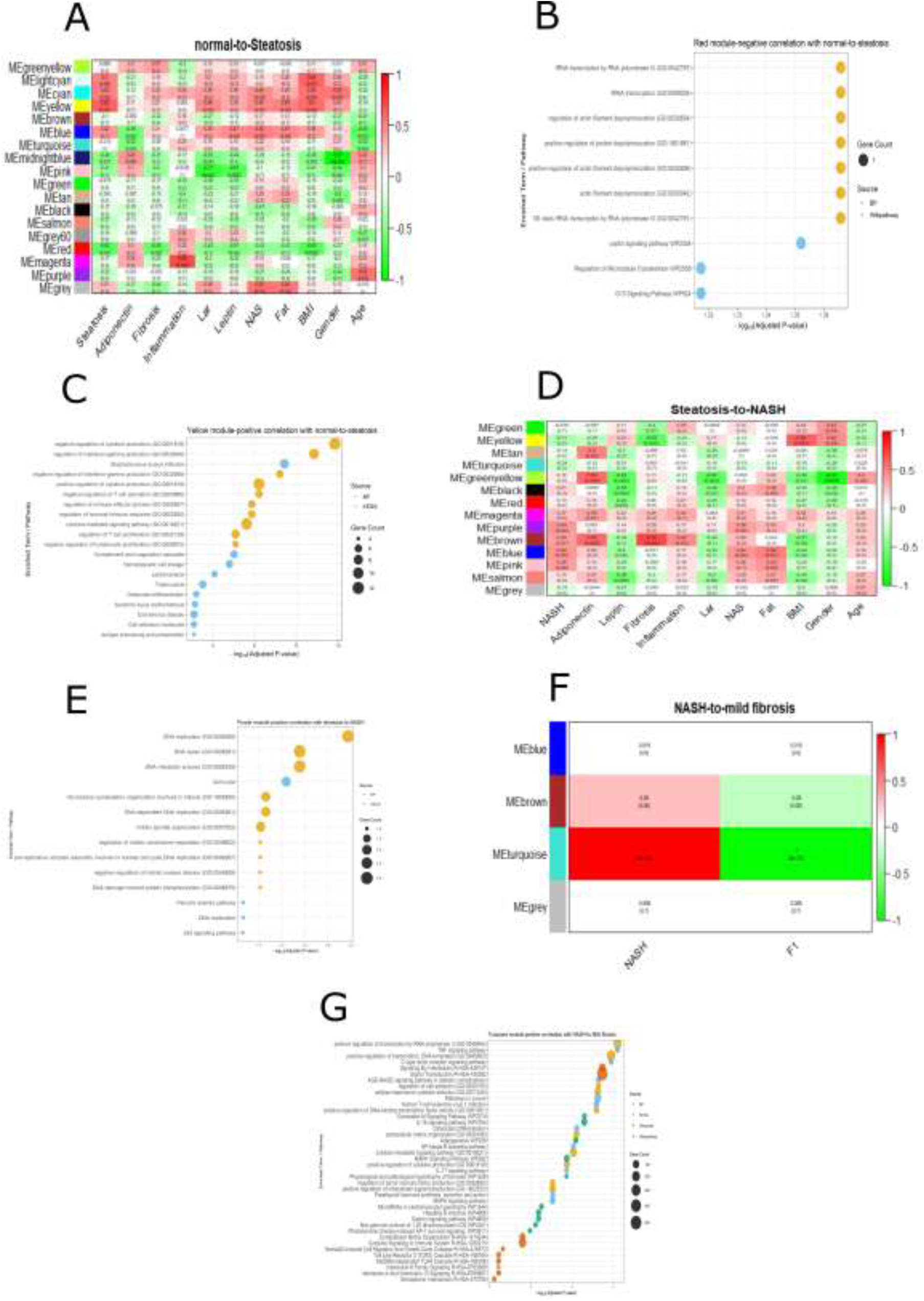
Weighted gene co-expression network analysis (WGCNA) identifies stage-associated co-expression modules. **a** Module–trait correlation heatmap for the normal→steatosis transition; each cell shows the correlation between a module eigengene and a clinical trait, with the correlation coefficient and *P* value (in parentheses). Red and green denote positive and negative correlations, respectively. **b** Functional enrichment of the Red module, negatively correlated with the normal→steatosis transition. **c** Functional enrichment of the Yellow module, positively correlated with the normal→steatosis transition. **d** Module–trait correlation heatmap for the steatosis→NASH transition. **e** Functional enrichment of the Purple module, positively correlated with the steatosis→NASH transition. **f** Module–trait correlation heatmap for the NASH→mild fibrosis transition. **g** Functional enrichment of the Turquoise module, correlated with the NASH→mild fibrosis transition. In enrichment panels, dot size represents gene count and colour represents the annotation source (BP, KEGG, Reactome, or WikiPathway); the *x* axis shows −log_10_(adjusted *P* value)

In the steatosis-to-NASH comparison, the Purple module was positively associated with disease-stage progression (r = 0.45, adjusted P = 0.04; Figure 3D). Functional enrichment analysis identified DNA replication and cell-cycle pathways among its enriched biological processes (Figure 3E). Although most hub genes in this module were not differentially expressed between steatosis and NASH, TCF19 and WDHD1 retained their hub-gene status despite relatively stable expression levels, suggesting that co-expression analysis may identify genes with important network roles even when their expression levels do not change significantly.

In the NASH-to-mild fibrosis comparison, the Turquoise module showed an inverse association with disease-stage progression (|r| > 0.25, adjusted P < 0.05; Figure 3F). This module was enriched in cell adhesion and TNF signaling pathways (Figure 3G).

In contrast, the mild-to-advanced fibrosis analysis did not identify any module with a statistically significant association with disease-stage progression.

Hub gene analysis identified PRDM1 and PLEK in the Yellow module, GTF3C6 and ZNF644 in the Red module, and TCF19 and WDHD1 in the Purple module (Tables S1 and S2). PRDM1 and PLEK exhibited dynamic expression changes during later disease stages, whereas GTF3C6 and ZNF644 showed altered expression patterns during fibrosis progression. TCF19 and WDHD1 met the predefined hub-gene criteria (|GS| > 0.2 and |MM| > 0.8), despite relatively stable expression levels. Overall, these findings highlight stage-associated co-expression patterns involving genes related to transcriptional regulation, cell proliferation, and genomic maintenance during NAFLD progression.

### Stage-Specific Remodeling of Protein–Protein Interaction Networks During NAFLD Progression

PPI network analysis revealed distinct, stage-dependent patterns of molecular interactions across NAFLD progression.

During the normal-to-steatosis transition, the PPI network comprised 31 nodes and 125 edges and was divided into two clusters (Figure 4A). Cluster 1, characterized by CXCR4 as its seed gene, was enriched in chemokine signaling, and all genes within this cluster showed downregulated expression (Figure 4B). Cluster 2 showed no significant functional enrichment (data not shown). These findings indicate changes in chemokine-associated interactions during the early transition from normal liver to steatosis.

**Figure 4.**
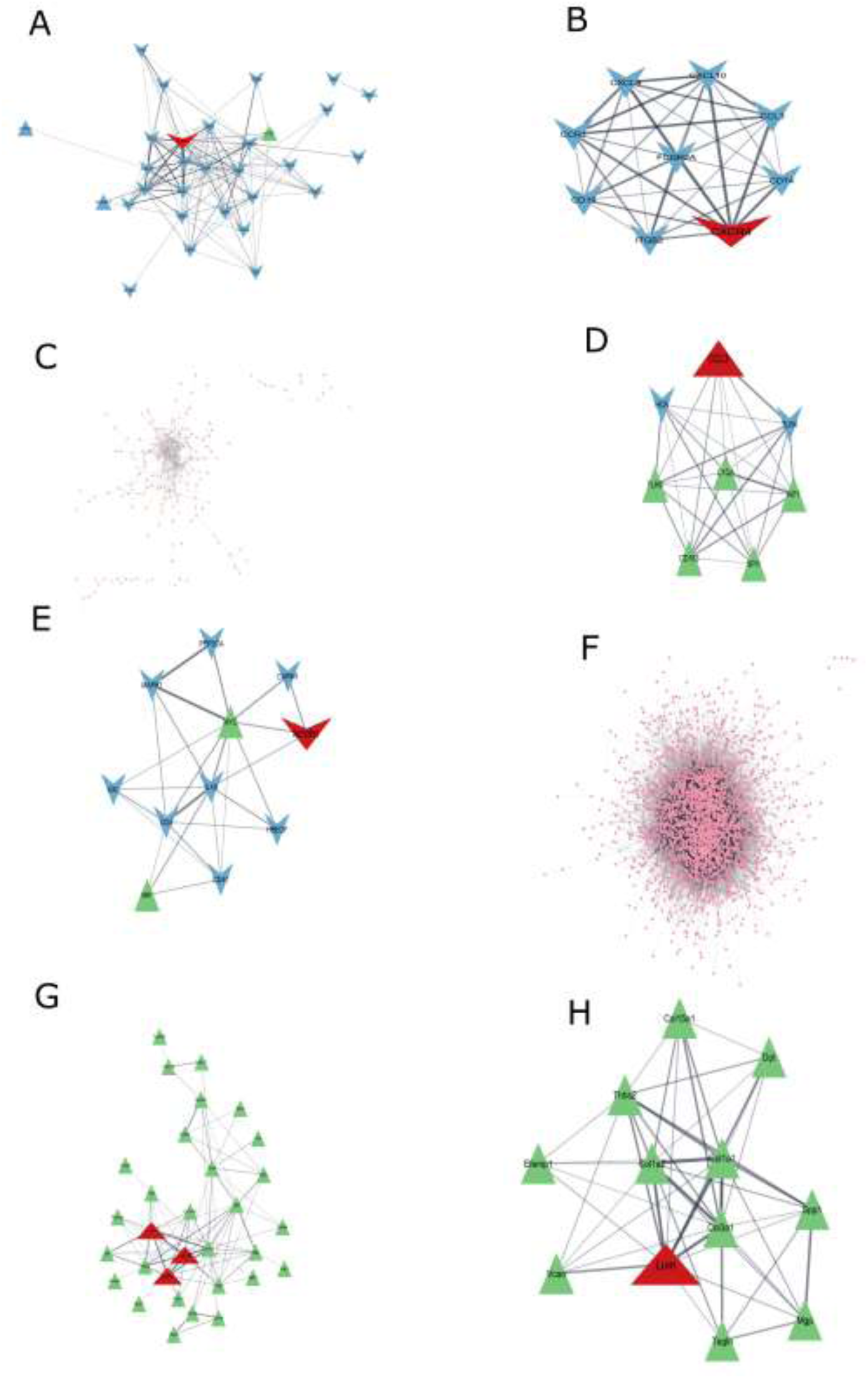
Stage-dependent remodeling of protein–protein interaction (PPI) networks across NAFLD progression. (A) Global PPI network for the normal-to-steatosis transition, comprising 31 nodes and 125 edges. (B) CXCR4-centered cluster from the normal-to-steatosis network, enriched in chemokine signaling, with all cluster genes showing downregulated expression. (C) Global PPI network for the steatosis-to-NASH transition, comprising 295 nodes and 541 edges. (D) CCL3-seeded cluster from the steatosis-to-NASH network, enriched in Toll-like receptor signaling. (E) NCOR1-seeded cluster from the steatosis-to-NASH network, enriched in ErbB signaling. (F) Global PPI network for the NASH-to-mild fibrosis transition, comprising 13 MCODE clusters. Detailed cluster information and functional enrichment results are provided in Supplementary File S2 and Table 2. (G) Global PPI network for the mild-to-advanced fibrosis transition. (H) LUM-associated Cluster 1 from the mild-to-advanced fibrosis network.

**Table 2.** Protein–Protein Interaction (PPI) Network Clusters Identified During the NASH-to-Mild Fibrosis Transition.

| **Functional Category** | **Cluster** | **Seed Gene** | **Key Enrichment / Pathway** |
|---|---|---|---|
| *Immune & Signaling* | 1 | CCL5 | PI3K-Akt & Chemokine signaling |
|  | 2 | CCL2 | Th17 cell differentiation |
|  | 6 | CLEC4E | SNARE interactions in vesicular transport |
| *Transcription & RNA Processing* | 4 | SKP2 | Spliceosome |
|  | 7 | COPA | Basal transcription factors |
|  | 8 | H1F0 | RNA transport |
| *Protein Homeostasis & Metabolism* | 10 | – | Ubiquitin-mediated proteolysis |
|  | 12 | – | Insulin resistance |
| *Ribosomal & Glycosylation* | 3 | TNFSF10 | Ribosome |
|  | 9 | SRGN | Glycosaminoglycan biosynthesis |
| *Unassigned / No Enrichment* | 5 | – | – |
|  | 13 | MICU2 | – |

During the steatosis-to-NASH transition, the PPI network expanded to 295 nodes and 541 edges (Figure 4C). Eight seed genes were identified: TMF1, PRPF8, ALG3, NCOR1, BPNT1, RGS1, CD69, and CCL3. Cluster 1, characterized by CCL3, was enriched in Toll-like receptor signaling, highlighting the involvement of innate immune and inflammatory pathways. Cluster 2, characterized by NCOR1, showed an overall downregulated expression pattern and was enriched in ErbB signaling (Figure 4D-E). These findings indicate changes in immune-related and growth factor-associated interactions during the transition from steatosis to NASH.

The NASH-to-mild fibrosis PPI network contained 13 MCODE clusters with distinct functional enrichment profiles (Figure 4F, Supplementary File S2). Enriched pathways included immune and signaling processes, such as PI3K-Akt and chemokine signaling (CCL5-associated Cluster 1), Th17 cell differentiation (CCL2-associated Cluster 2), and SNARE interactions in vesicular transport (CLEC4E-associated Cluster 6). Transcriptional and RNA-processing pathways included the spliceosome (SKP2-associated Cluster 4), basal transcription factors (COPA-associated Cluster 7), and RNA transport (H1F0-associated Cluster 8). Additional clusters were associated with ribosomal function (TNFSF10-associated Cluster 3), glycosaminoglycan biosynthesis (SRGN-associated Cluster 9), ubiquitin-mediated proteolysis (Cluster 10), and insulin resistance (Cluster 12).

During the mild-to-advanced fibrosis transition, one MCODE cluster was identified, with LUM as its seed gene (Figure 4G-H). Overall, these findings demonstrate stage-dependent changes in PPI network organization and functional enrichment across NAFLD progression, involving immune signaling, transcriptional regulation, RNA processing, and metabolic pathways.

### Validation of Steatosis Induction and Transgene Expression in HepG2 Cells

Fatty-acid treatment resulted in an approximately twofold increase in intracellular triglyceride levels compared with untreated cells (P < 0.01). Steatosis induction was further supported by increased expression of the lipogenic markers SREBP-1c, FASN, CD36, and PLIN2 (P < 0.01). Western blot analysis confirmed the expression of Myc-tagged CDKN1A and c-MYC proteins and detected EGFP as an independent marker of transfection (Figure 5A–D).

**Fig. 5.**
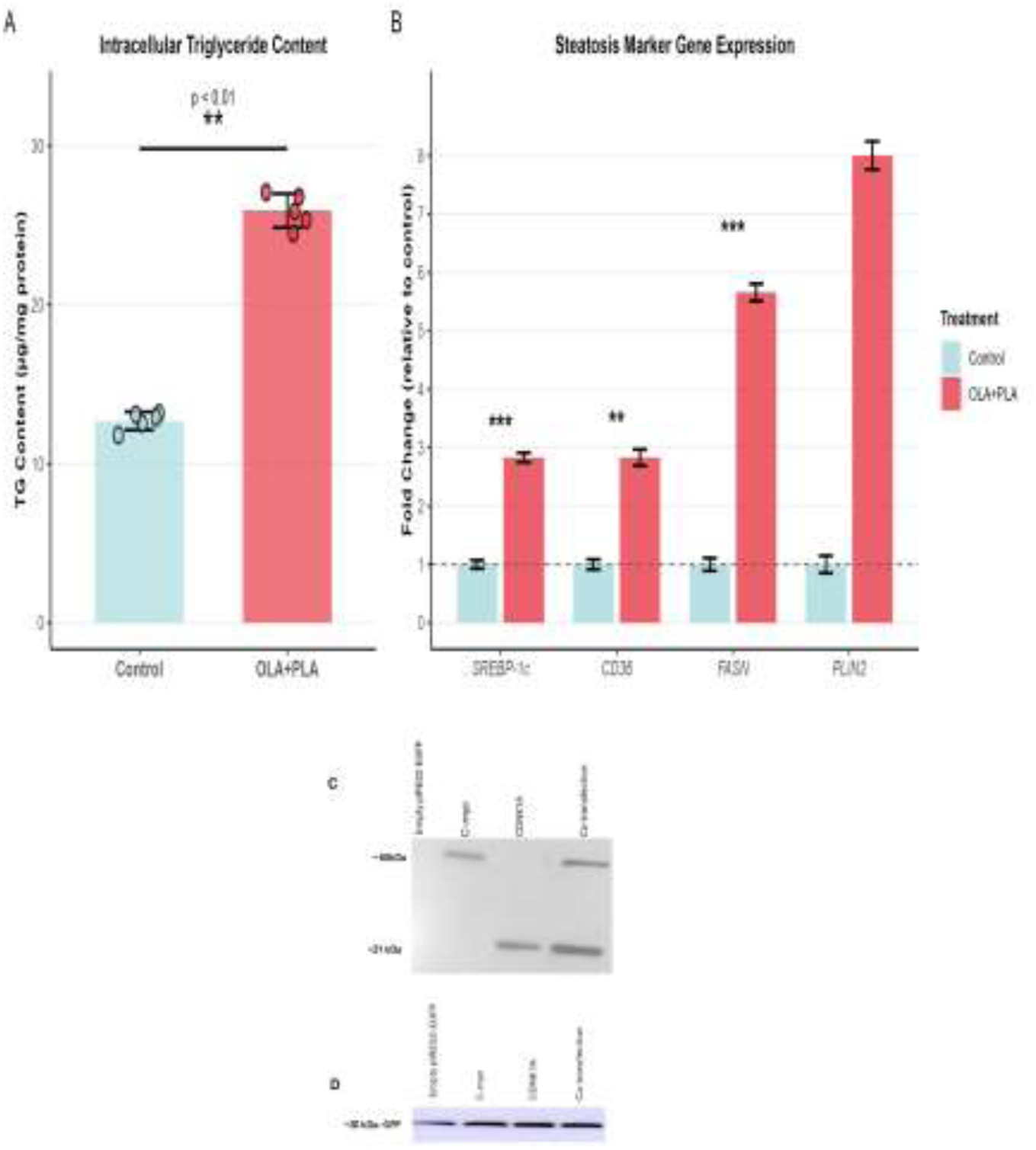
Induction of steatosis and validation of CDKN1A and c-MYC overexpression in HepG2 cells. **a** Intracellular triglyceride content in control and oleic acid/palmitic acid (OLA+PLA)-treated HepG2 cells (\*\**P* < 0.01). **b** Relative mRNA expression of steatosis-marker genes (*SREBP-1c*, *CD36*, *FASN*, *PLIN2*) in control and steatotic cells, expressed as fold change relative to control (\*\**P* < 0.01, \*\*\**P* < 0.001). **c** Western blot of myc-tagged proteins detected with an anti-myc-tag antibody: c-Myc (∼60 kDa) and CDKN1A/p21 (∼21 kDa) in empty-vector, c-Myc, CDKN1A, and co-transfection conditions. **d** Corresponding EGFP signal (∼27 kDa) confirming comparable transfection across conditions. Data are mean ± SD

### CDKN1A Overexpression Is Associated with Increased Inflammatory Gene Expression

The expression of selected genes involved in inflammation, immunity, metabolism, cell-cycle regulation, and TNF signaling was evaluated 72 hours after transfection (Figure 6). In steatotic HepG2 cells, CDKN1A overexpression was associated with increased expression of several inflammatory genes relative to the corresponding empty-vector control, including CCL20 (8-fold), IL6 (10-fold), CCL2 (10-fold), CCL3 (10-fold), and TNF (7-fold). In normal cells, the corresponding increases were generally smaller, ranging from approximately 2-to 4-fold for these genes.

**Figure 6.**
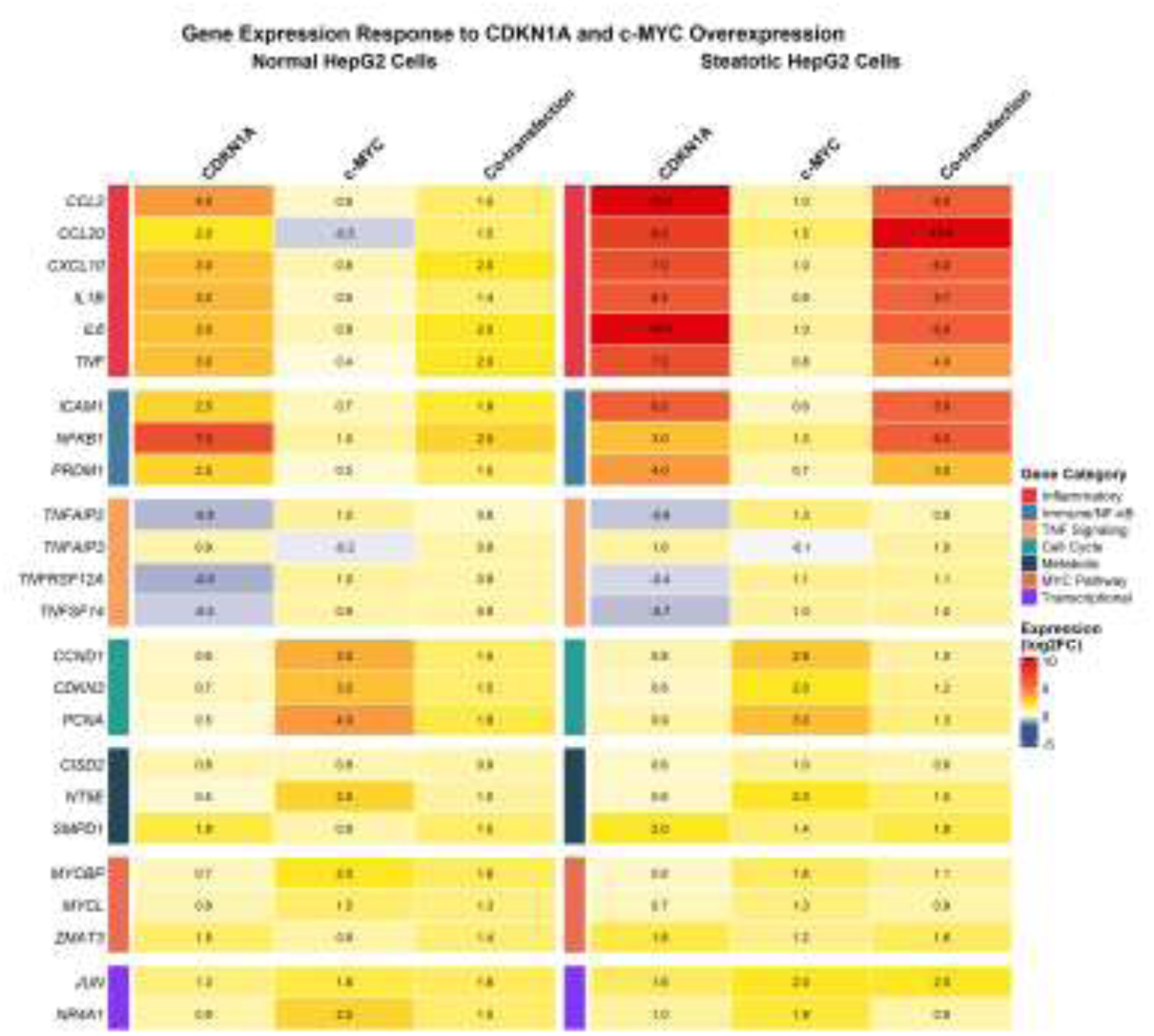
Expression profiles of 24 selected genes following CDKN1A and c-MYC overexpression in normal and steatotic HepG2 cells. Gene expression was assessed 72 hours after transfection in normal and fatty-acid-treated HepG2 cells. The 24 selected genes were grouped into seven functional categories: inflammatory genes (CCL20, IL6, TNF, CCL2, CXCL10, and IL1B); immune/NF-κB-related genes (ICAM1, NFKB1, and PRDM1); TNF-related genes (TNFAIP2, TNFAIP3, TNFRSF12A, and TNFSF14); cell-cycle-related genes (CDKN3, CCND1, and PCNA); metabolic genes (NT5E, CISD2, and SMPD1); MYC-pathway genes (MYCL, MYCBP, and ZMAT3); and transcriptional regulators (NR4A1, and JUN). Relative gene expression was determined using quantitative PCR and calculated using the 2⁻ΔΔCt method, with B2M as the reference gene. The CDKN1A, c-MYC, and co-transfection groups were compared with the corresponding empty-vector control within each cell condition. Data are presented as mean ± SEM from five independent biological replicates per group, with each sample analyzed in triplicate technical reactions.

In comparison, c-MYC overexpression was associated with relatively modest changes in the expression of these inflammatory genes in both cell conditions. Co-transfection produced gene-specific expression patterns: CCL20 and IL6 remained elevated (10-fold and 6-fold, respectively, in steatotic cells), whereas CCL2 and TNF showed lower expression than with CDKN1A overexpression alone.

CDKN1A overexpression was also associated with increased ICAM1 and NFKB1 expression in steatotic cells (6-fold and 3-fold, respectively). In normal cells, the corresponding values were 2.5-fold for ICAM1 and 7-fold for NFKB1. These findings indicate that CDKN1A overexpression was associated with changes in inflammatory gene expression in both cell conditions, although the magnitude and direction of the responses varied among genes.

### TNF-Related Genes Exhibit Differential Responses to CDKN1A and c-MYC Overexpression

The expression of four TNF-related genes, TNFAIP2, TNFAIP3 (A20), TNFRSF12A (Fn14), and TNFSF14 (LIGHT), was examined in normal and steatotic HepG2 cells following CDKN1A overexpression, c-MYC overexpression, or their co-transfection (Figure 6).

TNFAIP2 exhibited opposite expression patterns following CDKN1A and c-MYC overexpression. CDKN1A overexpression was associated with reduced TNFAIP2 expression in both normal and steatotic cells (log₂ fold change: −0.78 ± 0.15 and −0.60 ± 0.12, respectively). In contrast, c-MYC overexpression increased TNFAIP2 expression (log₂ fold change: +1.00 ± 0.12 in normal cells and +1.32 ± 0.11 in steatotic cells). Co-transfection resulted in intermediate expression levels (+0.58 ± 0.13 and +0.77 ± 0.12, respectively).

TNFAIP3 (A20) showed a different expression pattern. CDKN1A overexpression increased its expression in normal and steatotic cells (log₂ fold change: +0.87 ± 0.14 and +0.97 ± 0.12, respectively), whereas c-MYC overexpression was associated with modest reductions (−0.23 ± 0.14 and −0.10 ± 0.12, respectively). Following co-transfection, TNFAIP3 expression remained elevated (+0.90 ± 0.13 in normal cells and +1.00 ± 0.11 in steatotic cells).

A similar opposing pattern was observed for TNFRSF12A (Fn14). CDKN1A overexpression reduced its expression in both normal and steatotic cells (log₂ fold change: −0.87 ± 0.15 and −0.43 ± 0.14, respectively), whereas c-MYC overexpression increased expression (+1.00 ± 0.12 and +1.14 ± 0.12, respectively). Co-transfection was associated with increased TNFRSF12A expression (+0.90 ± 0.13 in normal cells and +1.10 ± 0.12 in steatotic cells).

TNFSF14 (LIGHT) also showed reduced expression following CDKN1A overexpression (log₂ fold change: −0.54 ± 0.12 in normal cells and −0.71 ± 0.14 in steatotic cells). In contrast, c-MYC overexpression increased TNFSF14 expression (+0.90 ± 0.12 and +1.00 ± 0.12, respectively). Co-transfection maintained increased expression in both conditions (+0.90 ± 0.11 and +1.00 ± 0.12, respectively).

Overall, these findings demonstrate distinct expression patterns among TNF-related genes following CDKN1A and c-MYC overexpression. CDKN1A was associated with increased TNFAIP3 expression and reduced TNFAIP2, TNFRSF12A, and TNFSF14 expression, whereas c-MYC generally showed the opposite pattern. Co-transfection produced gene-specific expression profiles that varied across the four genes.

### c-MYC Overexpression Is Associated with Increased Expression of Cell-Cycle-Related Genes

The expression of three cell-cycle-related genes, CDKN3, CCND1, and PCNA, was evaluated following CDKN1A overexpression, c-MYC overexpression, and CDKN1A/c-MYC co-transfection in normal and steatotic HepG2 cells (Figure 6).

c-MYC overexpression was associated with increased expression of all three genes in both cell conditions. CDKN3 expression increased approximately 3-fold in normal cells and 2-fold in steatotic cells. Similarly, CCND1 expression increased 3.5-fold in normal cells and 2.8-fold in steatotic cells, while PCNA expression increased 4-fold and 3-fold, respectively.

In contrast, CDKN1A overexpression was associated with relatively modest changes in these genes. CDKN3 expression remained below 1-fold in both conditions, whereas CCND1 and PCNA expression showed similarly limited changes. Co-transfection resulted in intermediate expression levels compared with the individual overexpression conditions. CDKN3 expression increased 1.5-fold in both normal and steatotic cells, while CCND1 expression increased 1.5-fold in normal cells and 1-fold in steatotic cells. PCNA expression increased 1.8-fold in normal cells and 1.3-fold in steatotic cells.

Overall, c-MYC overexpression was associated with higher expression of CDKN3, CCND1, and PCNA than CDKN1A overexpression in both normal and steatotic HepG2 cells. These findings indicate differential regulation of cell-cycle-associated gene expression by the two transcriptional regulators.

### Differential Expression of Metabolic and Transcriptional Regulators

The expression of genes involved in metabolic regulation and transcriptional control was also evaluated following CDKN1A overexpression, c-MYC overexpression, and their co-transfection in normal and steatotic HepG2 cells (Figure 6).

Among the metabolic regulators, CISD2 and SMPD1 showed relatively modest changes in expression across the experimental conditions. CISD2 expression remained close to baseline, with fold changes ranging from 0.6 to 1.0 across the different transfection conditions. SMPD1 showed more variable responses, with expression increases of up to 2-fold following CDKN1A overexpression in both normal and steatotic cells. NT5E (CD73) showed a greater response to c-MYC overexpression, with expression increasing 2.5-fold in normal cells and 2-fold in steatotic cells. Co-transfection resulted in more moderate increases (1-fold and 1.5-fold, respectively).

Among the transcriptional regulators, RORα and PRDM1 showed increased expression following CDKN1A overexpression, particularly in steatotic cells. RORα expression increased 4.3-fold in steatotic cells and 1.5-fold in normal cells, whereas PRDM1 expression increased 4-fold and 2.5-fold, respectively. In comparison, c-MYC overexpression was associated with relatively modest changes in both genes. Co-transfection resulted in intermediate expression levels.

NR4A1 showed relatively modest changes across the experimental conditions, with expression values generally ranging from 0.9-to 2.5-fold. HNF4α expression remained relatively stable, with values ranging from 0.5-to 1-fold across the different conditions.

Overall, these findings indicate gene-specific responses among metabolic and transcriptional regulators following CDKN1A and c-MYC overexpression, with particularly notable changes in NT5E, RORα, and PRDM1.

Expression of MYC-Associated Regulatory Genes Following CDKN1A and c-MYC Overexpression

The expression of three MYC-associated regulatory genes, MYCL, MYCBP, and ZMAT3, was examined to assess changes in the MYC-related transcriptional network following CDKN1A and c-MYC overexpression in normal and steatotic HepG2 cells (Figure 6).

c-MYC overexpression was associated with increased MYCL expression in both normal and steatotic cells (1.5-fold and 1.3-fold, respectively). MYCBP expression increased 2-fold in normal cells and 1.6-fold in steatotic cells. ZMAT3 expression showed a different pattern, increasing 0.9-fold in normal cells and 1.2-fold in steatotic cells following c-MYC overexpression.

CDKN1A overexpression was associated with relatively modest changes in these genes. MYCL expression was 0.8-fold in normal cells and 0.7-fold in steatotic cells, while MYCBP expression was 0.7-fold and 0.6-fold, respectively. ZMAT3 expression increased 1.5-fold in normal cells and 1.8-fold in steatotic cells.

Co-transfection produced intermediate expression patterns for MYCL and MYCBP, with fold changes of 1.3 and 1.6 in normal cells and 0.9 and 1.1 in steatotic cells, respectively. ZMAT3 expression was 1.4-fold in normal cells and 1.6-fold in steatotic cells.

Overall, MYCL, MYCBP, and ZMAT3 exhibited distinct expression responses to CDKN1A and c-MYC overexpression, with differences in the magnitude and direction of changes across the two cell conditions.

## DISCUSSION

### Stage-Specific Transcriptional Remodeling Defines NAFLD Progression

Our integrative analysis indicates that NAFLD progression is characterized by distinct, stage-dependent transcriptional changes rather than a uniform or linear molecular response. The number of DEGs varied substantially across consecutive disease-stage transitions, increasing from 34 in the normal-to-steatosis transition to 368 in the steatosis-to-NASH transition and reaching 3,208 during progression from NASH to mild fibrosis. In contrast, only 52 DEGs were identified between mild and advanced fibrosis. These findings suggest that the extent of transcriptional remodeling differs considerably across disease stages, with particularly extensive changes occurring during the transition from NASH to early fibrosis. This observation is consistent with previous studies demonstrating substantial cellular and molecular heterogeneity during NAFLD progression, including heterogeneous hepatocyte populations(Payen et al., 2021).

Further evidence of dynamic transcriptional regulation was provided by the 105 DEGs shared between the steatosis-to-NASH and NASH-to-mild fibrosis transitions, of which 46 exhibited inverse expression patterns between the two transitions. These genes were predominantly associated with inflammatory and NF-κB-related pathways, suggesting that inflammatory signaling undergoes dynamic changes during disease progression rather than following a consistently increasing or decreasing pattern. Such stage-dependent expression patterns may reflect changes in the biological responses of liver tissue as steatosis progresses toward steatohepatitis and fibrosis.

### Early Changes in Chemokine-Associated Signaling During Steatosis

An interesting finding of our study was the relatively limited transcriptional response during the normal-to-steatosis transition, accompanied by reduced expression of genes associated with chemokine signaling and immune-related processes. In particular, the CXCR4-centered PPI cluster showed enrichment in chemokine signaling, with downregulated expression of all genes within the cluster. These findings suggest that early hepatic steatosis may involve changes in immune-associated signaling that do not necessarily correspond to an immediate increase in inflammatory gene expression.

Although NAFLD is commonly characterized by progressive inflammatory changes, the molecular response during its early stages may be more complex than a simple increase in inflammatory activity(Arrese, Cabrera, Kalergis, & Feldstein, 2016). Reduced expression of selected chemokine-associated genes could reflect an early adaptation to metabolic stress or changes in the regulation of immune-cell recruitment and communication within the liver(Fasano et al., 2022; Wang et al., 2023). However, because our analysis was based on bulk liver transcriptomic data, it cannot establish whether these changes originate from hepatocytes, resident immune cells, or alterations in the proportions of different hepatic cell populations.

Importantly, the subsequent increase in differentially expressed genes during the steatosis-to-NASH transition, together with the emergence of inflammatory and Toll-like receptor-associated interactions in the PPI network, suggests a substantial shift in the molecular environment as steatosis progresses toward steatohepatitis. Further studies using single-cell or spatial transcriptomic approaches will be necessary to determine the cellular sources and functional consequences of these early changes in chemokine-associated signaling.

### WGCNA Reveals Latent Regulatory Programs

WGCNA identified stage-associated co-expression modules involving genes related to inflammatory signaling, cell-cycle regulation, and tissue remodeling during NAFLD progression. Notably, TCF19, WDHD1, GTF3C6, and ZNF644 were identified as hub genes despite relatively stable expression levels, highlighting the importance of network connectivity beyond individual gene-expression changes. Previous studies have implicated TCF19 and WDHD1 in cellular proliferation and genomic maintenance, suggesting their potential relevance to hepatocyte responses during disease progression(Li & Wang, 2024; Y. Liu et al., 2025; Vasilopoulos & Martínez-Zamudio, 2024). Furthermore, the enrichment of complement and coagulation pathways in the Yellow module suggests a possible link between immune and vascular-associated processes during early steatosis. These findings highlight potential regulatory pathways that warrant further functional investigation.

### CDKN1A as a Context-Dependent Inflammatory Regulator

Our experimental findings suggest that CDKN1A overexpression is associated with changes in inflammatory gene expression in HepG2 cells, with response patterns varying according to the metabolic condition of the cells. In steatotic cells, CDKN1A overexpression was associated with increased expression of several inflammatory mediators, including CCL20, IL6, CCL2, CCL3, TNF, CXCL10, and IL1B. These changes were also observed in normal cells, although their magnitude differed among genes and experimental conditions. This pattern suggests that the inflammatory gene-expression response associated with CDKN1A may be influenced by the metabolic state of hepatocytes.

Previous studies have suggested that p21 may participate in cellular stress responses and inflammatory regulation through interactions with signaling pathways and transcriptional regulators, including NF-κB, activator protein 1, and signal transducers and activators of transcription(Elmitwalli et al., 2024; Lukosz et al., 2010; Stojanovic et al., 2025; Q. Yu et al., 2025). Metabolic stress and oxidative stress have also been implicated in the regulation of p21 expression and activity. These mechanisms may provide a biological context for the differences observed between normal and steatotic HepG2 cells. However, the present study did not directly investigate p21 protein localization, its interactions with transcription factors, or the activation status of these signaling pathways.

An additional finding was the increased expression of TNFAIP3, which encodes the anti-inflammatory regulator A20, following CDKN1A overexpression in both normal and steatotic cells. A20 is known to participate in the negative regulation of inflammatory signaling, particularly through its effects on NF-κB-associated pathways. Its increased expression alongside several inflammatory mediators suggests that CDKN1A-associated responses may involve both pro-inflammatory and potentially counter-regulatory components. Nevertheless, the functional contribution of A20 to these responses remains to be established.

Taken together, these findings suggest that CDKN1A is associated with a context-dependent inflammatory gene-expression profile in HepG2 cells. Its effects appear to involve changes in multiple inflammatory and regulatory genes rather than a uniform increase in inflammatory signaling. Further studies examining p21 protein activity, NF-κB signaling, cytokine production, and A20 function are needed to clarify the mechanisms underlying these observations.

### c-MYC Is Associated with Increased Expression of Cell-Cycle-Related Genes

In contrast to the inflammatory gene-expression changes associated with CDKN1A overexpression, c-MYC overexpression was associated with increased expression of several cell-cycle-related genes in both normal and steatotic HepG2 cells. In particular, CDKN3, CCND1, and PCNA showed increased expression following c-MYC overexpression, with changes observed under both metabolic conditions. These findings suggest that c-MYC may contribute to the regulation of cell-cycle-associated transcriptional programs in hepatocytes, independently of whether the cells are exposed to fatty acids.

The expression patterns of other MYC-associated genes, including MYCL, MYCBP, and ZMAT3, were more variable, indicating that the transcriptional response to c-MYC overexpression was not uniform across the MYC-related gene network. These differences may reflect gene-specific regulatory mechanisms and interactions between c-MYC-associated pathways and the metabolic state of the cells.

The role of c-MYC in cell growth and proliferation has been extensively investigated, and dysregulated c-MYC signaling has been associated with tumor development and hepatocellular carcinoma. Previous experimental studies have also demonstrated that sustained c-MYC overexpression can contribute to liver injury, steatohepatitis, fibrosis, and hepatocarcinogenesis in transgenic models(Hou et al., 2017; Jia, Che, Cigliano, Wang, Peitta, Tao, Zhong, Ribback, Evert, Chen, et al., 2020; F. Liu, Liao, & Zhang, 2023). In addition, the increased expression of NT5E observed following c-MYC overexpression is noteworthy because NT5E encodes CD73, an enzyme involved in extracellular adenosine production. Adenosine signaling has been implicated in the regulation of immune responses and tumor progression(Nevzorova et al., 2016).

Although these findings provide a potential link between c-MYC-associated transcriptional changes, cell-cycle regulation, and metabolic signaling, our experiments did not directly assess cell proliferation, cell-cycle distribution, or tumor-related phenotypes. Therefore, the observed changes in gene expression should be interpreted as evidence of altered cell-cycle-associated transcription rather than direct evidence of increased proliferation.

### The CDKN1A-c-MYC Axis Is Associated with Differential Expression of TNF-Related Genes

An additional finding of this study was the distinct expression patterns of four TNF-related genes—TNFAIP2, TNFAIP3, TNFRSF12A, and TNFSF14—following CDKN1A and c-MYC overexpression. CDKN1A overexpression was associated with reduced expression of TNFAIP2, TNFRSF12A, and TNFSF14, accompanied by increased TNFAIP3 expression. In contrast, c-MYC overexpression was associated with increased expression of TNFAIP2, TNFRSF12A, and TNFSF14, while TNFAIP3 showed relatively modest reductions. Co-transfection produced gene-specific expression patterns, indicating that the combined expression of these regulators was not associated with a uniform response across the four genes.

These findings are particularly relevant to TNFRSF12A (Fn14), which encodes the receptor for TNF-like weak inducer of apoptosis (TWEAK). TWEAK–Fn14 signaling has been implicated in inflammatory responses, hepatocyte injury, and activation of hepatic fibrogenic processes(Y. Zhang, Zeng, & Xia, 2021). Previous studies have also reported associations between increased TWEAK signaling and the severity of steatohepatitis(Dertli et al., 2019). The increased TNFRSF12A expression observed following c-MYC overexpression therefore raises the possibility that c-MYC-associated transcriptional changes may influence pathways involved in hepatic inflammation and tissue remodeling. However, the present study did not directly assess Fn14 protein expression or TWEAK-mediated signaling.

Similarly, TNFSF14 encodes LIGHT, a member of the TNF superfamily that has been associated with inflammatory signaling and cellular responses, including cytokine production and apoptosis(Otterdal et al., 2015). Its increased expression following c-MYC overexpression suggests a potential relationship between c-MYC-associated transcriptional regulation and LIGHT-related inflammatory pathways. Previous work has also suggested interactions between LIGHT-associated signaling and inflammatory mediators(J. Zhang et al., 2023). Nevertheless, whether these interactions contribute to the expression patterns observed in our experimental model remains unknown.

Overall, the differential expression of TNFAIP2, TNFAIP3, TNFRSF12A, and TNFSF14 highlights the involvement of TNF-related regulatory genes in the transcriptional responses associated with CDKN1A and c-MYC overexpression. These findings suggest that the CDKN1A-c-MYC axis may be relevant to the regulation of inflammatory and tissue-remodeling pathways during NAFLD progression. Further experiments examining protein expression, receptor signaling, and functional interactions between these pathways are required to determine whether these transcriptional changes translate into altered TNF-related signaling activity.

### Therapeutic Implications and Stage-Specific Research Directions

The stage-dependent molecular changes identified in this study suggest that therapeutic strategies for NAFLD may benefit from considering the distinct biological processes associated with different stages of disease progression. The relatively limited transcriptional changes observed during early steatosis, followed by more extensive inflammatory and fibrotic remodeling during subsequent stages, highlight the importance of understanding how metabolic stress and inflammatory responses evolve over time.

During early disease, interventions targeting metabolic dysfunction may help limit progression toward steatohepatitis. Therapeutic approaches involving glucagon-like peptide-1 receptor agonists and sodium-glucose cotransporter-2 inhibitors have attracted attention because of their metabolic effects and potential benefits in patients with metabolic dysfunction-associated liver disease(Rajewski et al., 2025). Although these treatments were not evaluated in the present study, the early transcriptional changes identified here provide a rationale for further investigating the relationship between metabolic regulation and the initiation of inflammatory responses.

The CDKN1A-c-MYC axis may also represent an interesting area for future investigation. The association of CDKN1A overexpression with inflammatory gene expression, together with the increased expression of cell-cycle-related genes following c-MYC overexpression, suggests that these regulators may contribute to distinct aspects of the hepatic response to metabolic stress. In particular, the role of p21-associated cellular senescence and the senescence-associated secretory phenotype warrants further investigation. Although senolytic approaches have shown promise in experimental models of chronic liver disease, their potential effects may depend on disease stage and the biological functions of senescent cells(David S Umbaugh, Diehl, & Du, 2025).

Similarly, c-MYC-associated transcriptional changes may warrant further investigation in the context of hepatocyte proliferation and liver tumor development. However, because c-MYC has important physiological functions, therapeutic targeting would require careful evaluation of efficacy and potential effects on normal liver homeostasis(F. Liu et al., 2023). The differential expression of TNFRSF12A and TNFSF14 also highlights TWEAK–Fn14 and LIGHT-associated pathways as potential areas for further study, particularly in relation to hepatic inflammation and fibrotic remodeling(Romero-Suárez, Solana-Tinoco, García-Espiñeira, Lambis-Anaya, & Suarez-Causado, 2024). Finally, the increased expression of TNFAIP3 following CDKN1A overexpression raises the possibility that A20-associated regulatory mechanisms may contribute to counter-regulatory responses during inflammatory stress. Further investigation of TNFAIP3 and its functional effects may help clarify its potential relevance to liver disease(da Silva et al., 2025; D. Liu et al., 2020).

Overall, these findings provide a basis for investigating stage-specific molecular targets in NAFLD. However, additional studies using primary hepatocytes, multicellular liver models, and in vivo experimental systems are necessary before the therapeutic relevance of the CDKN1A-c-MYC axis and associated inflammatory pathways can be established.

### Study Limitations

This study has several limitations. First, the use of bulk liver transcriptomic datasets limits the identification of cell-specific expression changes and does not distinguish transcriptional alterations from changes in cellular composition. Second, the cross-sectional nature of the datasets limits causal and temporal interpretations of NAFLD progression. Third, HepG2 cells cannot fully reproduce the complexity of the human liver, and the experimental findings require validation in primary hepatocytes and more physiologically relevant models. Furthermore, our experiments focused on gene expression and did not directly assess protein function, inflammatory signaling, cell proliferation, or senescence. Finally, WGCNA and PPI analyses identify associations and predicted interactions rather than experimentally confirmed regulatory mechanisms. Further functional and multi-omics studies are needed to validate these findings.

### Conclusions

This study provides an integrated overview of stage-associated molecular changes during NAFLD progression, identifying distinct transcriptional profiles, co-expression networks, and protein–protein interaction patterns across disease stages. The findings indicate that early steatosis is associated with relatively limited transcriptional remodeling and changes in chemokine-associated signaling, followed by more extensive inflammatory and molecular changes during the transition to NASH and subsequent fibrotic progression.

Experimental findings further suggest that CDKN1A and c-MYC overexpression are associated with distinct, context-dependent gene-expression responses in normal and steatotic HepG2 cells. CDKN1A overexpression was associated with changes in inflammatory and TNF-related genes, whereas c-MYC overexpression was associated with increased expression of several cell-cycle-related genes. These observations highlight the potential relevance of the CDKN1A-c-MYC regulatory axis to the transcriptional responses of hepatocytes under metabolic stress.

Although further functional validation is required to establish the underlying mechanisms, the integration of transcriptomic, network-based, and experimental analyses provides a framework for investigating stage-associated molecular pathways and potential regulatory targets during NAFLD progression.

## ACKNOWLEDGMENTS

The authors thank the investigators who made their expression datasets publicly available through the Gene Expression Omnibus. We also thank Shiraz University of Medical Sciences for financial support through PhD thesis grant no. 98-01-74-21114 awarded to Z.M.

## CRediT Author Contributions

Zahra Moayedfard: Conceptualization, Methodology, Investigation, Data curation, Formal analysis, Writing – original draft.

Negar Azarpira: Conceptualization, Methodology, Investigation, Data curation, Formal analysis, Writing – original draft.

Shima Parsa: Conceptualization, Writing – review & editing. Kimia Falamarzi: Conceptualization, Writing – review & editing. Farnaz Sani: Conceptualization, Writing – review & editing.

Ayeh Bolouki: Conceptualization, Methodology, Investigation, Data curation, Formal analysis, Supervision, Project administration, Writing – review & editing.

All authors reviewed and approved the final manuscript and agree to be accountable for its contents.

## Data Availability Statement

The data supporting the findings of this study are publicly available in the Gene Expression Omnibus (GEO) database (https://www.ncbi.nlm.nih.gov/geo/). The analyses were based on the following datasets: GSE48452 (https://www.ncbi.nlm.nih.gov/geo/query/acc.cgi?acc=GSE48452), GSE49541 (https://www.ncbi.nlm.nih.gov/geo/query/acc.cgi?acc=GSE49541), and GSE63067 (https://www.ncbi.nlm.nih.gov/geo/query/acc.cgi?acc=GSE63067).

Processed data, analysis code, and supplementary materials are available at: https://github.com/AyehBlk/NAFLD-transcriptomics-analysis

## Ethics Approval Statement

This study analyzed publicly available, de-identified gene expression datasets. Ethical approval for the original studies was obtained from the respective institutions, as reported in the original publications (Ahrens et al., 2013; Murphy et al., 2013; Frades et al., 2015). No new human participants were recruited for this study.

## Informed Consent Statement

Not applicable.

## Consent for Publication

Not applicable.

---

**Table S1.** Top 10 hub genes ranked by GS in modules with a significant correlation between GS and MM.

| **Normal → Steatosis** |  |  |  | **Steatosis → NASH** |  | **NASH → Mild fibrosis** |  |
|---|---|---|---|---|---|---|---|
| **Red** |  | **Yellow** |  | **Purple** |  | **Turquoise** |  |
| **Gene** | **GS** | **Gene** | **GS** | **Gene** | **GS** | **Gene** | **GS** |
| **RBM7** | -0.68 | **AXL** | 0.84 | **CHEK1** | 0.53 | **HSPA6** | 0.999875 |
| **ZNF644** | -0.68 | **CSF1R** | 0.82 | **KNTC1** | 0.51 | **PAX8** | 0.999816 |
| **CFL2** | -0.66 | **CR1** | 0.75 | **WDHD1** | 0.47 | **UBA7** | 0.999802 |
| **ZNF875** | 0.65 | **CD4** | 0.74 | **FANCI** | 0.44 | **MAPK1** | 0.999782 |
| **GTF3C6** | -0.63 | **SDC3** | 0.73 | **MCM6** | 0.44 | **PRR22** | 0.999756 |
| **MORN2** | -0.63 | **FCGR2B** | 0.71 | **MLF1IP** | 0.43 | **SLC39A13** | 0.999695 |
| **SKA2** | -0.62 | **CD163** | 0.70 | **CENPK** | 0.42 | **TRIOBP** | 0.999651 |
| **MRPL42** | -0.62 | **CD180** | 0.69 | **TCF19** | 0.41 | **TVP23C** | 0.999634 |
| **NPBWR1** | 0.61 | **MPEG1** | 0.69 |  |  | **TSSK3** | 0.999626 |
| **HSF4** | 0.60 | **CD74** | 0.69 |  |  | **DSCR10** | 0.999605 |

**Table S2.**
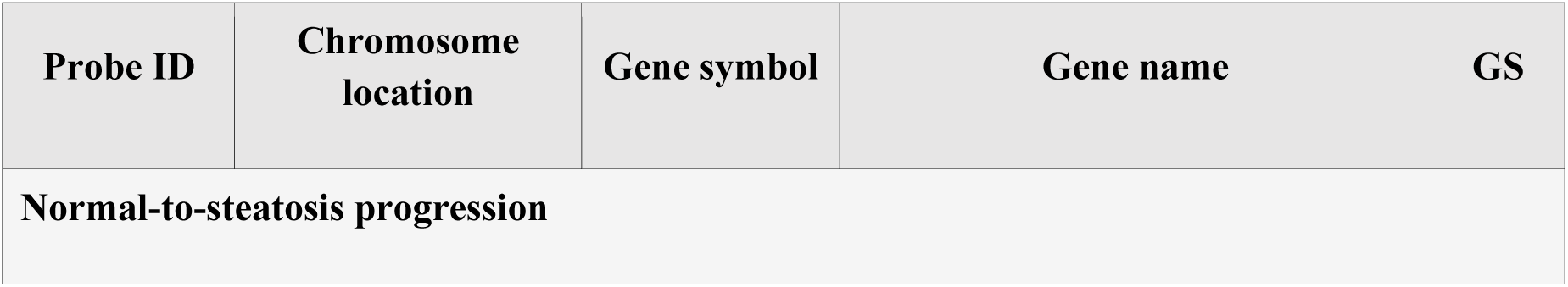
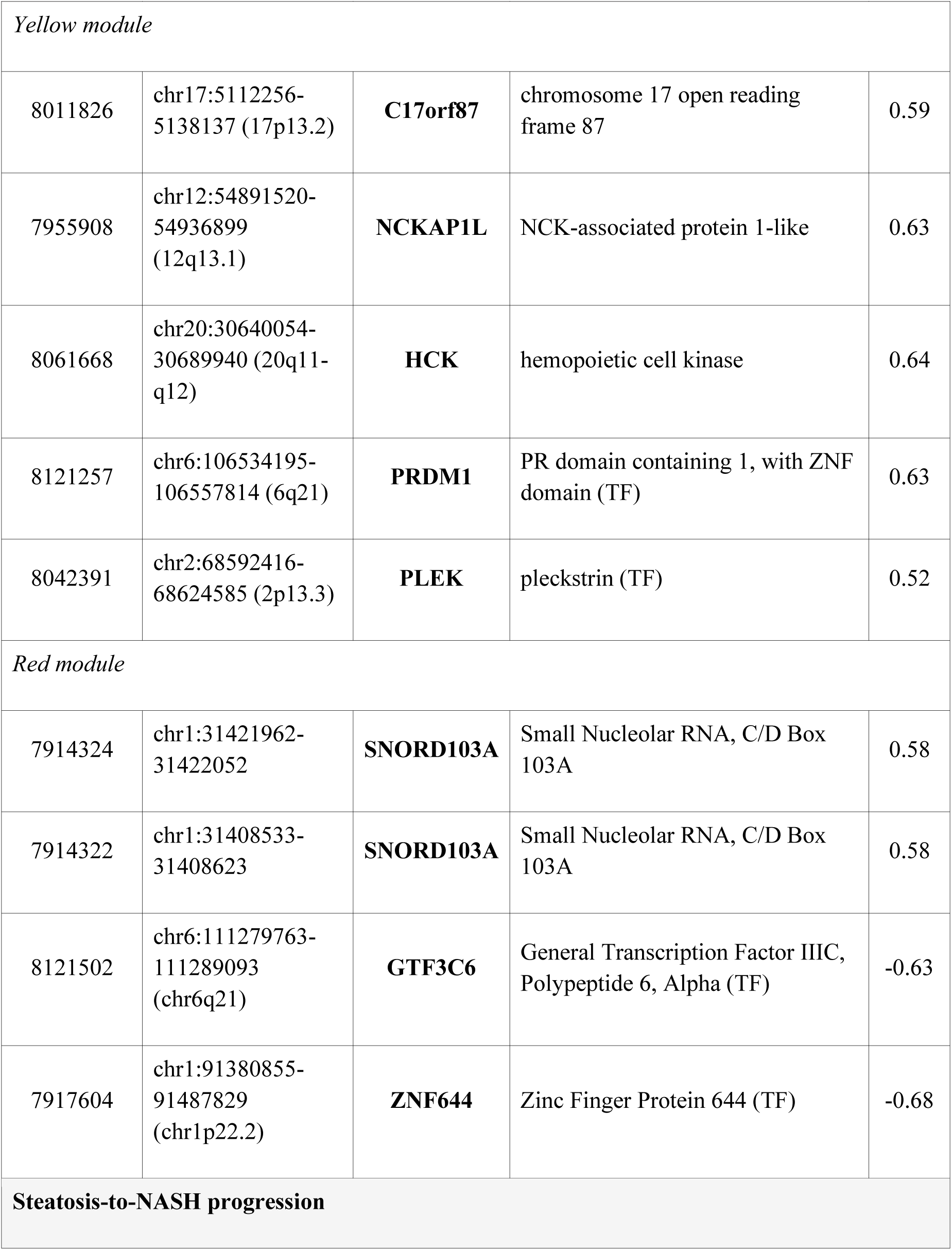
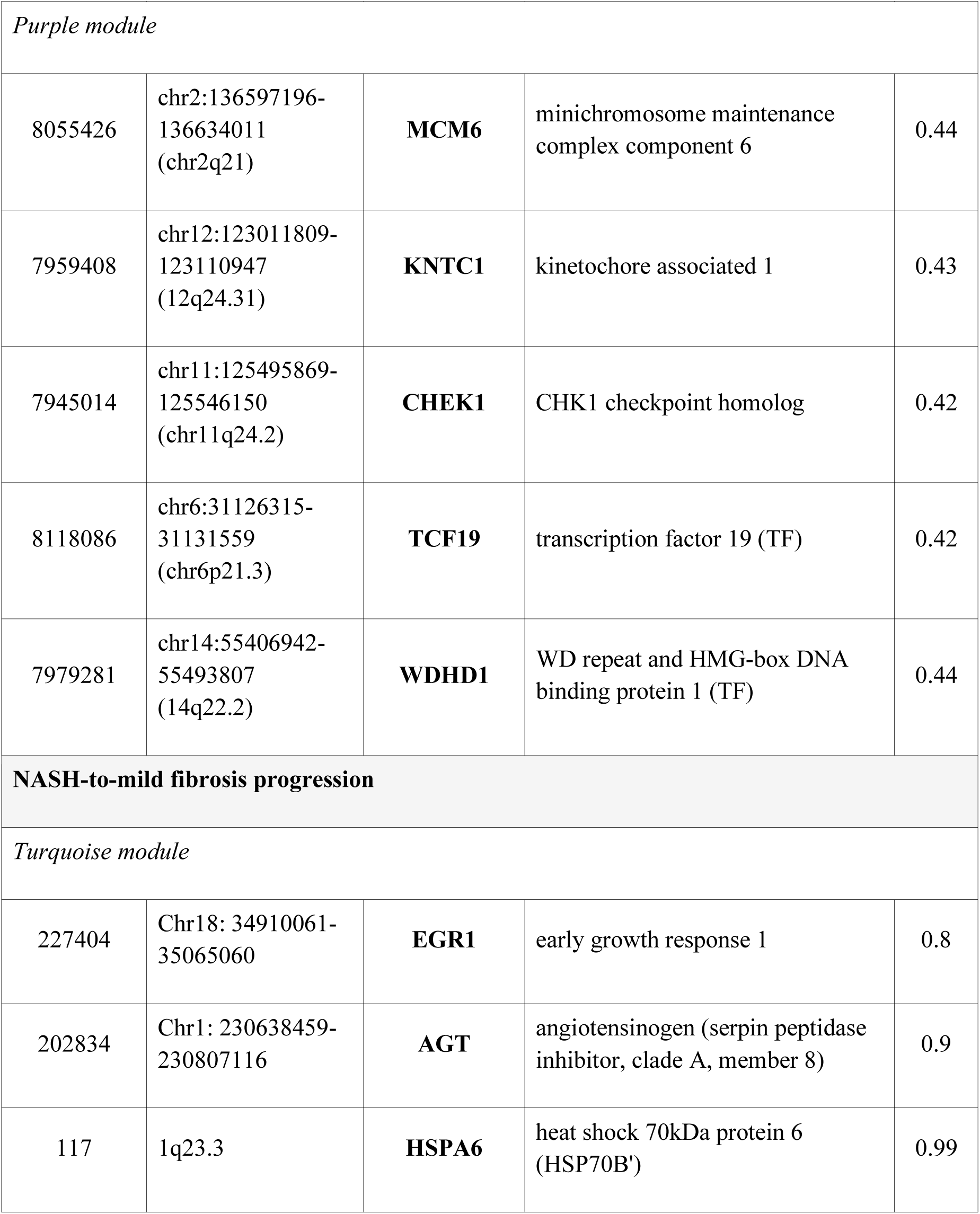
TFs and hub nodes in the module co-expression networks determined by weight and degree scores.

| Probe ID | Chromosome location | Gene symbol | Gene name | GS |
|---|---|---|---|---|
| **Normal-to-steatosis progression** |  |  |  |  |
| *Yellow module* |  |  |  |  |
| 8011826 | chr17:5112256-5138137 (17p13.2) | **C17orf87** | chromosome 17 open reading frame 87 | 0.59 |
| 7955908 | chr12:54891520-54936899 (12q13.1) | **NCKAP1L** | NCK-associated protein 1-like | 0.63 |
| 8061668 | chr20:30640054-30689940 (20q11-q12) | **HCK** | hemopoietic cell kinase | 0.64 |
| 8121257 | chr6:106534195-106557814 (6q21) | **PRDM1** | PR domain containing 1, with ZNF domain (TF) | 0.63 |
| 8042391 | chr2:68592416-68624585 (2p13.3) | **PLEK** | pleckstrin (TF) | 0.52 |
| *Red module* |  |  |  |  |
| 7914324 | chr1:31421962-31422052 | **SNORD103A** | Small Nucleolar RNA, C/D Box 103A | 0.58 |
| 7914322 | chr1:31408533-31408623 | **SNORD103A** | Small Nucleolar RNA, C/D Box 103A | 0.58 |
| 8121502 | chr6:111279763-111289093 (chr6q21) | **GTF3C6** | General Transcription Factor IIIC, Polypeptide 6, Alpha (TF) | -0.63 |
| 7917604 | chr1:91380855-91487829 (chr1p22.2) | **ZNF644** | Zinc Finger Protein 644 (TF) | -0.68 |
| **Steatosis-to-NASH progression** |  |  |  |  |
| *Purple module* |  |  |  |  |
| 8055426 | chr2:136597196-136634011 (chr2q21) | **MCM6** | minichromosome maintenance complex component 6 | 0.44 |
| 7959408 | chr12:123011809-123110947 (12q24.31) | **KNTC1** | kinetochore associated 1 | 0.43 |
| 7945014 | chr11:125495869-125546150 (chr11q24.2) | **CHEK1** | CHK1 checkpoint homolog | 0.42 |
| 8118086 | chr6:31126315-31131559 (chr6p21.3) | **TCF19** | transcription factor 19 (TF) | 0.42 |
| 7979281 | chr14:55406942-55493807 (14q22.2) | **WDHD1** | WD repeat and HMG-box DNA binding protein 1 (TF) | 0.44 |
| **NASH-to-mild fibrosis progression** |  |  |  |  |
| *Turquoise module* |  |  |  |  |
| 227404 | Chr18: 34910061-35065060 | **EGR1** | early growth response 1 | 0.8 |
| 202834 | Chr1: 230638459-230807116 | **AGT** | angiotensinogen (serpin peptidase inhibitor, clade A, member 8) | 0.9 |
| 117 | 1q23.3 | **HSPA6** | heat shock 70kDa protein 6 (HSP70B') | 0.99 |

